# Structural and Synaptic Dysregulation Drives Network Vulnerability in APOE4 Human Neuronal Networks

**DOI:** 10.64898/2026.05.04.722197

**Authors:** Marthe Bendiksvoll Grønlie, Vegard Fiskum, Axel Sandvig, Ioanna Sandvig

## Abstract

Synaptic failure and associated neuronal network dysfunction are key pathological processes involved in the early stages of Alzheimer’s disease (AD). A better understanding of the specific synaptic pathways and network topologies that drive disease vulnerability is therefore essential for the development of a targeted therapeutic intervention. In the present study, we aimed to determine how defined synaptic pathways and connectivity patterns shape the emergence and progression of the structural and functional network dynamics of human neuronal networks with inherent vulnerability to AD. We performed longitudinal microelectrode array recordings, assessed excitatory and inhibitory activity, quantified neurite growth, and performed proteomic analyses of synaptosomes from human induced pluripotent stem cell-derived neuronal networks carrying homozygous apolipoprotein E epsilon 4 (APOE4), the strongest genetic risk factor for developing late-onset AD. This integrated approach enabled multiscale characterization of synaptic alterations, structural maturation, and functional network dynamics associated with AD vulnerability. Compared to isogenic homozygous APOE3 networks, we found that APOE4 drives a distinct topological regime, characterized by high assortativity combined with low transitivity, which reflects a compensatory organization with reduced redundancy and flexibility, consistent with an intrinsically fragile network structure. APOE4 networks exhibited reduced firing rates, dynamic excitatory and inhibitory imbalance, impaired synchronization, absence of network bursting, and reduced global routing efficiency. Despite retaining small-world properties indicative of baseline information processing capacity, the topological and functional profile of APOE4 networks suggests a reliance on compensatory mechanisms associated with elevated metabolic cost and increased susceptibility to pathological spread. Structurally, APOE4 networks displayed reduced dendritic length, branching, and total dendrite area, accompanied by dysregulation of synaptic organization and signaling, ion dynamics, and intracellular signaling pathways. Together, these findings establish that APOE4 drives a multiscale reorganization of neuronal networks that not only mirrors synaptic alterations identified in patients, but also contextualizes these changes within network-level dynamics, advancing a more comprehensive understanding of early AD pathology.

## Introduction

Neurodegenerative diseases are characterized by progressive and selective neuronal dysfunction and cell death, often preceded by synaptic and network-level disturbances (Kvistad et al., 2024) (Ming and Song, 2011). A prominent manifestation of such dysfunction is an imbalance between excitatory and inhibitory synaptic transmission, which has been increasingly recognized as a shared pathological feature across Alzheimer’s disease (AD), Parkinson’s disease, and amyotrophic lateral sclerosis (Tanaka et al., 2025). Network vulnerability presents differently across disorders, reflecting the selective susceptibility of distinct neuronal populations and brain circuits. In AD, the most prevalent neurodegenerative disease, early pathological changes preferentially affect memory-related circuits within the medial temporal lobe, and subsequently involve distributed cortical regions. (Braak and Braak, 1991) (Thal et al., 2002) (Zhang et al., 2024) (de Wilde et al., 2016).

Network-level dysfunction is consistently associated with hallmark AD neuropathologies, including amyloid-β (Aβ) accumulation (Hardy and Allsop, 1991) (Hardy and Higgins, 1992) (Gallego Villarejo et al., 2022) (Caramello et al., 2025) and neurofibrillary tangles (Braak and Braak, 1991) (Thal et al., 2002). Although the mechanistic and temporal relationships between Aβ and tau pathology remain unresolved (Kepp et al., 2023), growing evidence suggests that neuronal dysfunction disrupts synaptic communication and consequently alters emergent network dynamics across spatial scales. Importantly, synaptic dysfunction in AD is increasingly recognized as an early and progressive systems-level disturbance rather than merely a secondary consequence of overt neuronal loss (Burns et al., 2026) (Li et al., 2025) (Palop and Mucke, 2010) (Castanho et al., 2025). Neuroimaging studies have demonstrated altered task-related activation of hippocampal and cortical brain regions, years before clinical diagnosis both in presymptomatic carriers of AD-related mutations (Quiroz et al., 2010) (Ereira et al., 2024) (Petrella et al., 2011), and in healthy individuals with genetic risk (Filippini et al., 2009) (Bookheimer et al., 2000). Complementary findings from transgenic models show aberrant functional connectivity even in the absence of substantial Aβ deposition or overt synaptic loss (Morrissey et al., 2023) (Mondragón-Rodríguez et al., 2019) (Zerbi et al., 2014). Together, these findings indicate that altered network dynamics is an early feature of AD.

The increased prevalence of epileptiform activity in AD patients, often emerging prior to overt cognitive decline (Ranasinghe et al., 2025), suggests an early pathological phenotype characterized by network hyperexcitability. How-ever, altered E/I balance in AD cannot be reduced to simple runaway excitation. Instead, accumulating evidence shows that pathological processes differentially affect excitatory and inhibitory circuits across brain regions and disease stages, resulting in dynamic and context-dependent network states (Burns et al., 2026) (Li et al., 2025). In early stages of the disease, heightened excitatory drive may recruit compensatory inhibitory responses, temporarily stabilizing network activity. Such homeostatic plasticity can support initial functional resilience. However, as the pathological burden increases with disease progression and age, the brain’s compensatory capacity is exceeded, leading to persistent network instability and progressive dysfunction (Palop and Mucke, 2010) (Castanho et al., 2025) (De Strooper and Karran, 2016).

At the level of large-scale network organization, functional connectivity findings are similarly heterogeneous, with reports of both hyper- and hypoconnectivity across disease stages (Zerbi et al., 2014) (Fadel et al., 2025). Cellular models recapitulate this variability, showing altered firing patterns and synchrony without a consistent electrophysiological signature (von Maydell et al., 2025) (Ghatak et al., 2024) (Ortiz-Virumbrales et al., 2017) (Kawatani et al., 2024) (Ghatak et al., 2021). Rather than reflecting inconsistency, this variability suggests that AD-related network dysfunction represents a dynamic and context-dependent process.

Despite extensive evidence that AD involve early synaptic and network-level dysfunction, it remains to be unraveled how these alterations evolve over time to shape the emergent architecture of human neuronal networks. In particular, few studies have examined how intrinsic vulnerability to AD influences the longitudinal neuronal network activity and connectivity at high temporal resolution. Moreover, the relationship between altered network dynamics and underlying synaptic function remains largely unresolved. In the current study, we aimed to determine how defined synaptic pathways and connectivity patterns shape the emergence and progression of the structural and functional network dynamics of human neuronal networks with inherent vulnerability to AD.

We performed longitudinal microelectrode array recordings to capture the spontaneous network activity and connectivity of human neuronal networks with genetic vulnerability to AD, i.e. carrying two alleles of the APOE4 gene. Although both APOE4 and isogenic APOE3 control networks possessed information processing capacity, the efficiency of information flow and functional integration was hampered in APOE4 networks. APOE4 networks showed reduced firing rates compared to controls, which was compensated for by an increased fraction of spikes in electrode bursts and higher assortativity, the latter suggesting that nodes of similar degree tend to connect to each other. Higher connectivity of hub nodes protects important pathways from perturbations, however, at the cost of rendering the networks vulnerable to disease spread. The lower ratio of triangles to triples in APOE4 networks, transitivity, indicated an overall vulnerability to network failure, with few redundant pathways when comparing all pairs of nodes. Hence, the functional organization of APOE4 networks represent a compensatory and intrinsically fragile network topology. We also demonstrated a time-dependent shift in E/I balance in APOE4 networks, with early elevated excitatory drive transitioning to increased fraction of inhibitory synaptic connections at identified later time points. Underlying this functional vulnerability, synaptosome proteomics revealed a dysregulation of synaptic organization, synaptic signaling, ion dynamics, and intracellular signaling in APOE4 networks. Structurally, APOE4 neurons organized into networks with shorter and less branched dendrites, accompanied by dysregulation of actin, dendrite, dendritic tree, cell adhesion, and motility. Together, these structural, molecular, and functional alterations demonstrate that APOE4 does not simply suppress network activity, but reconfigures neuronal networks into a state that is simultaneously compensatory and inherently vulnerable to failure.

## Methods

### Differentiation of human cortical neurons

Human induced pluripotent stem cells (iPSCs) were purchased from The Jackson Laboratory. The JIPSC001142 cell line, carrying two APOE4 alleles, was used to model genetic vulnerability to AD (Safiri et al., 2024). To study APOE4-specific effects, the isogenic parental cell line KOLF2.1J (JIPSC001000, passage number 4, male, 55-59 yrs) harboring two APOE3 alleles was used as control (Pantazis et al., 2022). APOE3 is associated with a neutral risk of developing AD (Safiri et al., 2024). The iPSCs were expanded following the manufacturers instructions in StemFlex Medium (Gibco, A3349401) on 100 mm petri dishes (Corning, Falcon, 353003) coated with 5 ug/mL vitronectin (Gibco, A14700). When the cells reached 80% confluence, neural rosette differentiation was initiated following a protocol adapted from Dannert et al. (2023).

To promote cell viability, iPSCs were supplemented with 10 uM Y-27632 dihydrochloride (RI) (Sigma-Aldrich, Y0503) added to the cell medium 1h prior to passaging. 12-well plates (Corning, Costar, 3512) were coated with 100 ug/mL Geltrex (Gibco, A1413201) and incubated at 37°C, 5% CO2 for 1h and then at RT for 1h. After removing the old medium, cells were washed with Dulbecco’s Phosphate Buffered Saline without calcium and magnesium (PBS-/-) (Sigma-Aldrich, D8537) and incubated in 2 mL Accutase (Gibco, A1110501) at 37°C, 5% CO2 until individual cell detachment was observed under a microscope after approximately 10 min. To neutralize Accutase, 4 mL DMEM/F12 (Gibco, 11320074) supplemented with 10 uM RI was added before transferring the cells to a pre-rinsed conical tube. Cells were centrifuged at 170 x g for 4 min, and the supernatant was removed before resuspending the cell pellet in Neural Induction Medium (NI) (Dannert et al.,2023) supplemented with 10 uM RI. A total of 2 mL NI + RI containing 8 × 10^5^ cells were plated in each well of the Geltrex-coated 12-well plate. This time point was defined as 0 days in vitro (DIV) until this was redefined on the day of the final plating. Cell medium was replaced daily with 2 mL of fresh NI until the cells were passaged at 6 DIV.

6-well plates (Corning, Falcon, 353046) were coated with 50 ug/mL Poly-L-ornithine (PLO) (Sigma-Aldrich, P4957) in RT overnight. PLO was removed and replaced with 10 ug/mL laminin (Gibco, 23017015) for 2h at 37°C, 5% CO2. Laminin was removed and plates were left open to dry in the laminar flow hood for 30 min until an even crystalline sheet appeared. After removing the old medium, cells were washed with PBS-/- and incubated in 0.5 mL Accutase as described above. Following neutralization of the Accutase with 1 mL DMEM/F12 + RI, cells were centrifuged at 170 x g for 4 min. The pellet was resuspended in NI + RI and diluted to a final concentration of 24 M cells/mL. For spot plating, a 250-350 uL drop of the cell suspension was added per well of the dried 6-well plate. The drop was kept intact while very gently tilting the plate until the drop covered 1/3 to 1/2 of the well’s diameter. After placing the lid back on, cells were incubated at 37°C, 5% CO2 for 30 min to facilitate attachment before gently adding 2 mL NI + RI. The next day (7 DIV), the cell medium was replaced with 4 mL fresh NI. Starting at 9 DIV the medium was replaced with 4 mL Neural Differentiation Medium (NM) (Dannert et al., 2023) every two days onward. When neural rosettes appeared, around 15-20 DIV, 20 ng/mL Basic Fibroblast Growth Factor (bFGF) (Stemcell Technologies, 78003.1) was added to the cells for two subsequent days to support rosette proliferation and expansion while continuing regular NM media changes every two days. When the neural rosettes started accumulating and dense circles were visible under the microscope around 20-25 DIV, rosettes were passaged for further expansion.

6-well plates were coated with PLO and laminin as described above, but without leaving the plates open to dry. After removing the old medium from the cells, 1 mL STEMdiff Neural Rosette Selection Reagent (NRSR) (Stemcell Technologies, 05832) was added to each well followed by 45 min incubation at 37°C, 5% CO2. The NRSR was removed and replaced with 1.5 mL NM + RI. A micropipette tip was used to manually isolate the edges of the spot from its center. Cells from the center were transferred to a conical tube and triturated to break up clumps. Cells were centrifuged at 350 x g for 5 min, the supernatant was removed, and the pellet was resuspended in NM supplemented with 20 ng/mL bFGF and 10 uM RI. Cells in 2 mL medium were plated directly after liquid laminin was removed, at a 3:2 to 3:4 ratio depending on the size and density of the spots. The next day, the medium was replaced with fresh NM + bFGF, before subsequent feeds with NM only every two days onward. When the rosettes became almost confluent around 30-33 DIV, they were passaged as individual cells using Accutase.

6-well plates were coated with PLO and liquid laminin as described above. After removing the old medium, the cells were washed with PBS-/- and incubated with 0.5 mL Accutase for 20-30 min at 37°C, 5% CO2. After neutralizing the Accutase using 1 mL NM, the cells were transferred to a conical tube and centrifuged at 170 x g for 4 min, resuspended in NM + bFGF + RI and plated in 2 mL medium per well. Three confluent wells were passaged onto 3-5 new wells depending on their densities. The next day, the medium was replaced with fresh NM + bFGF, before subsequent feeds with NM only every two days onward.

When the cells had re-formed rosettes and reached 100% confluence, typically 4-7 days after Accutase split, they were passaged twice more at 4-7 day intervals, following the procedure described above. During the third and final passage, cells were resuspended in NB+ / B27+ (Dannert et al., 2023) supplemented with 10 uM DAPT (Selleck Chemicals, S2215) and 10 uM RI, and plated at a final density of 2000 cells/mm^2^. 0 DIV was now redefined as the day of the final plating. Cells were plated onto 8-well chamber slides (Nunc, LabTek, 177445) for immunocytochemistry, Cytoview 6-well MEAs (Axion Biosystems, M384-tMEA-6) for electrophysiological recordings, and 6-well plates (Corning, Falcon, 353046) for proteomics analyses. When plating cells onto the MEAs, 210 uL cell suspension was added onto the electrode areas only. The cells were then incubated for 30 min at 37°C, 5% CO2 to facilitate attachment before gently adding medium to obtain a final volume of 1 mL.

At 2 DIV, all the old medium was replaced with NB+ / B27+ supplemented with DAPT. At 3 DIV, 50% of old medium was replaced with NB+ / B27+ supplemented with DAPT, 5 uM 5-Fluorouracil (Sigma-Aldrich, F6627), and 5 uM Uridine (Sigma-Aldrich, U3750). 5-Fluorouracil and Uridine (collectively referred to as 5FU) were added to obtain a pure neuronal culture by removing proliferating cells. Subsequent media changes were performed every two days, with DAPT being added for a total of 7 days and 5FU for a total of 12 days. Starting at 15 DIV and onward, cells were maintained by replacing 50% of the old medium with fresh NB+ / B27+ every two days. The full experimental timeline from this point is illustrated in Fig. 1 and described below.

**Fig. 1.**
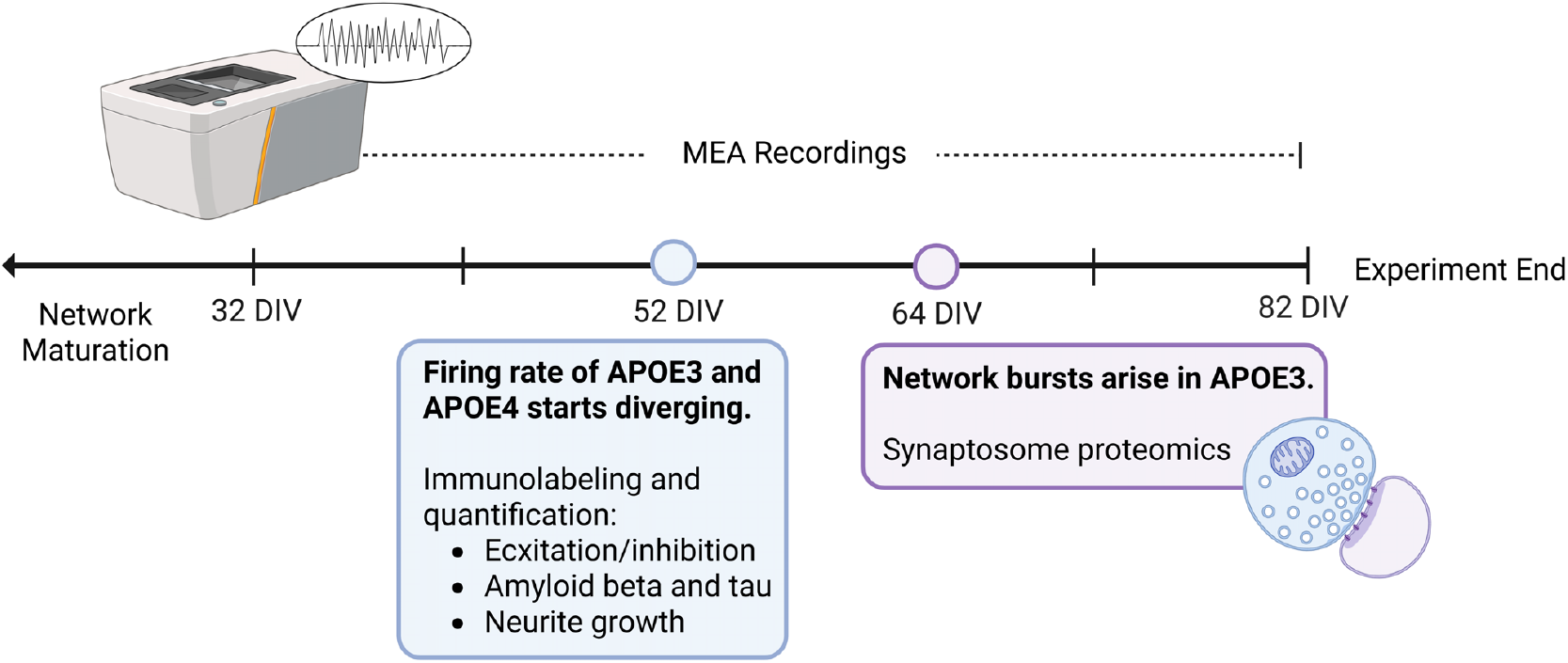
Human induced pluripotent stem cell-derived cortical neurons were matured on microelectrode arrays (MEAs) until spontaneous action potentials emerged. Twenty-minute recordings were initiated at the onset of individual spike activity (32 days in vitro (DIV)) and conducted every two days until 66 DIV, then every four days until 82 DIV. Sixty-minute recordings were additionally obtained from 70–82 DIV to enable analysis of excitatory and inhibitory network links. At 52 DIV, immunolabeling was performed to quantify excitatory and inhibitory neurotransmission, amyloid-β and tau pathology, and neurite morphology, in order to investigate mechanisms underlying diverging firing rates between genotypes. From 64 DIV onwards, network bursting was exclusive to APOE3; synaptosomes were therefore isolated at this time point for proteomic analysis to identify mechanisms driving this functional divergence.

### Electrophysiological recordings and analysis

Extra-cellular recordings of neural network activity were obtained using the Axion Maestro Pro MEA system (Axion Biosystems, GA, USA). The AxIS Navigator software version 3.10.3.6 was used for data acquisition at a sampling rate of 12.5 kHz. To allow network activity to stabilize after transport to the recording system, the cells were left to rest for 30 min at 37°C, 5% CO2 before each recoding. 20 min recordings were obtained from 32-68 DIV, before 60 min recordings were obtained from 70-82 DIV to enable analysis of excitatory and inhibitory links. Recordings were initiated when the first individual spikes were observed at 32 DIV, and subsequently obtained every two days until 66 DIV, followed by recordings every four days until 82 DIV. Spike and burst detection was performed in Axis Navigator. Spike detection was performed within a filtered frequency band of 300-3000 Hz using an adaptive threshold of 6 standard deviations from the continuous stream, a pre-spike duration of 0.84 ms and a post-spike duration of 2.16 ms. An electrode burst was detected when at least 5 spikes were detected within a maximum inter-spike interval (ISI) of 100 ms. A network burst was detected when at least 25% of the electrodes detected at least 50 spikes in total within a maximum ISI of 100 ms. The synchrony window was set to 20 ms.

Matlab (version R2023a) (MathWorks, 2023) was used to summarize the spike and burst statistics obtained from Axis Navigator. Recordings were excluded from the analysis if containing less than 7 spikes per min or less than 5 active electrodes. The Information Theory Toolbox (Timme and Lapish, 2018) was used to calculate functional connectivity based on mutual information. Graph theoretical metrics (Table 1) were calculated using the Brain Connectivity Toolbox (Rubinov and Sporns, 2010). Excitatory and inhibitory network edges were distinguished using the method outlined in Pastore et al. (2018). Briefly, network edges were identified by cross-correlation with a time window of 30 ms and distinguished as excitatory or inhibitory based on the sign of the cross-correlogram peak with the highest absolute value. The assessment of excitatory and inhibitory links was only performed on recordings obtained at 70 DIV and later time points because a valid analysis requires a sufficiently high firing rate.

**Table 1.** Graph Parameters and their definitions.

| Graph Parameter | Description |
| --- | --- |
| Global Routing Efficiency | Average inverse shortest path length across all pairs of nodes. Large values indicate efficient information flow and functional integration (Latora and Marchiori, 2001) (Goñi et al., 2013). |
| Small World Propensity | Combination of local clustering and average path length. High clustering and short path length, providing a value $> .6$ is considered small world, and support information processing capacity (Muldoon et al., 2016). |
| Assortativity | Correlation between node degrees on opposite sides of a link. Stronger correlation supports robust information transfer, but also increases the vulnerability to targeted damage (Foster et al., 2010) (Newman, 2002). |
| Transitivity | Ratio of triangles to triplets. Higher transitivity indicates more redundancy and functional flexibility (Rubinov and Sporns, 2010) (Onnela et al., 2005). |

### Statistical analysis of electrophysiology

SPPS (version 30.0.0.0 (172)) (IMB Corp, 2024) was used for statistical analysis of the electrophysiological data. Generalized linear mixed models were fitted to test if the neuronal activity slopes differed between APOE4 and APOE3 networks. Outliers were removed if located more than three inter-quartile ranges outside the middle 50% of the data without any clear pattern of group or time point. The distributions of the electrophysiology metrics were assessed by Shapiro Wilk test, frequency histograms, and Q-Q plots for each group. Considering that all electrophysiology metrics were continuous and that neuronal activity data is often long-tailed (Nobukawa, 2022) (Chow et al., 2025), the normal and gamma distributions were used for model fitting. In the case of normality, normality of Pearson residuals was confirmed after model fitting. For skewed electrophysiology distributions, gamma with log link was used as target distribution. The electrophysiology metrics were selected as targets, with the appropriate target distributions and link functions as indicated in Table 3. Model fit was evaluated based on the Akaike and Bayesian information criteria. Individual networks were selected as subjects, and time points were selected as repeated measures. First-order autoregressive (AR1) covariance structure was used to account for the dependent observations within wells. AR1 assumes that data points obtained closer in time are more correlated than those further apart. Group, time points, and group*time points including intercepts were selected as fixed effects. For all electrophysiology metrics except firing rate and small world propensity, the individual networks including intercepts were selected as random effects to model individual variability between networks and account for the possibility of different baselines between each network. For firing rate and small world propensity this random effect was excluded from the models because their random effects variance components were approximately 0, meaning that the random effect contained so little variance that it was redundant and caused convergence issues. Satterthwaite approximation was used to calculate effective degrees of freedom. Except from that, default build options in SPSS were used. Group differences were estimated using pairwise contrasts while keeping time points constant at their mean. Sequential Bonferroni was used to adjust for multiple comparisons. Results were considered significant if adjusted p<.05. Estimated means were displayed in terms of the original target scales. The data and output from Matlab and SPSS were imported to RStudio (RStudio IDE 2025.05.0+496) (R Core Team, 2021) using the packages haven (version 2.5.5) (Wickham et al., 2025b) and readxl (version 1.4.5) (Wickham and Bryan, 2025), and plotted using dplyr (version 1.1.4) (Wickham et al., 2025a), tidyr (version 1.3.1) (Wickham et al., 2025c), and ggplot2 (version 3.5.2) (Wickham, 2016).

### Synaptosome isolation

To investigate synaptic pathways underlying the functional network activity, synaptosomes were isolated from neurons on 6-well plates at 64 DIV using Syn-PER Synaptic Protein Extraction Reagent (Thermo Fisher, 87793) and EDTA-free Halt Protease and Phosphatase Inhibitor Cocktail (Thermo Fisher, 78441) according to the manufacturer’s instructions. Cells were washed twice with PBS-/- before Syn-PER application. Cells were scraped off and centrifuged at 1200 x g for 15 min, before the supernatant was transferred to a new tube. The supernatant was then centrifuged at 15,000 x g for 30 min, before the new supernatant was removed and discarded. The remaining synaptosome pellet was resuspended in Syn-PER with protease and phosphatase inhibitor and dimethyl sulfoxide (Sigma-Aldrich, D8418) and stored at -80 °C until analysis. All reagents were applied cold and the centrifugation steps were performed at 4 °C.

### Mass spectrometry

Mass spectrometry (MS) was conducted by the Norwegian University of Science and Technology (NTNU) Proteomics and Modomics Experimental Core Facility (PROMEC). Synaptosome preparations were resuspended in 1% sodium deoxycholate, 100mM Tris-hydrochloride (pH 8.5), 10 mM Tris(2-carboxyethyl)phosphine, and 40 mM chloroacetamide. Using a Bioruptor Pico instrument (Hologic Diagenode, B01080010), the samples were sonicated for 10 cycles of alternating 30 s sonication and 30 s rest. Following heating of the samples at 95°C for 30 min, they were added 0.01 M ammonium bicarbonate and 0.5 ug trypsin for digestion at 37°C overnight. Desalting of the resulting peptide suspension was performed using C18 spin columns. Using a SpeedVac vacuum concentrator centrifuge (Thermo Fisher), the peptides were dried and subsequently reconstituted in 0.1% formic acid. Liquid chromatography (LC) with tandem MS was performed using a timsTOF Pro with the nonoElute LC system (Bruker Daltonics). Peptide separation was achieved during a 75 min run using a 25 cm × 150 um column (Pepsep 25, Bruker Daltonics) with running buffers (A) 0.1% formic acid and (B) buffer A in acetonitrile (2-40% gradient). The timsTOF Pro was used in DIA PASEF mode with 1,600 V capillary voltage; 3,0 L/min dry gas; and 180°C dry temperature. MS1 and MS2 spectra were acquired within a 100-1,700 m/z range, with collision energy linearly interpolated between 59 eV (1/K0 = 1.60 V*·*s/cm^2^) and 20 eV (1/K0 = 0.6 V*·*s/cm^2^).

### Preprocessing of mass spectrometry data

The DIANN academic version 2.2.0 (Demichev et al., 2022) was used for processing raw MS data from the timsTOF Pro. Along with defaults, oxidation of methionine and acetylation of protein N-terminal as dynamic post-translational modification was appended. Both the MS and MS/MS mass-accuracy was set to 20 PPM. Search was performed against the library that is generated with human proteome consisting of one protein sequence per gene downloaded from Uniprot in December 2024 along with DIANN’s internal contaminants database. QuantUMS (high accuracy) quantification strategy (Kistner et al., 2023) was utilized to reduce missing values in the raw data, with protein inference enabled and match-between-runs (MBR) functionality. MBR verifies the presence of a peptide signal in one sample that was not identified in another. Precursor identification false discovery rate was kept at 1% and only those with higher confidence were used for the final protein group identification. Intensity values for each protein-group were normalized using DIANN’s implementation of label-free quantification algorithm (Cox et al., 2014). Quality control check was done using an in-house developed R script.

### Bioinformatic enrichment analysis

MS intensity report files were analyzed in RStudio (RStudio IDE 2025.09.2+218) (R Core Team, 2021). Differentially expressed proteins (DEPs) were identified and plotted using the DEP package (version 1.26.0) (Zhang et al., 2018). Only proteins identified in all samples from at least one condition were included in the analysis. Data was normalized by variance stabilizing transformation. Missing data patterns were evaluated by plotting missing data as a heatmap, and by plotting intensity distributions and the cumulative fractions of proteins with and without missing values. Left-censored missing data was imputed by random draws from a Gaussian distribution centered to the .01-th quantile of the observed values in each sample. Proteins were considered differentially expressed if they passed thresholds of p<.05 and log2 fold change ≥1.5. Functional enrichment analysis was performed using the clusterProfiler package (version 4.12.6) (Yu, 2024) (Xu et al., 2024) (Wu et al., 2021) (Yu et al., 2012). Gene ontology (GO) over-representation analysis (Boyle et al., 2004) was performed to identify which processed the most enriched proteins are involved in. GO over-representation analysis was performed on identified up- and downregulated DEPs, with Benjamini–Hochberg correction for multiple comparisons. Enriched GO terms were plotted using enrichplot (version 1.24.4) (Yu, 2023). Gene set enrichment analysis (GSEA) (Subramanian et al., 2005) was performed to identify processes that were more subtly up- or downregulated by considering trends in the total set of identified proteins. GSEA was performed with number of permutations set to 10,000; minimum and maximum gene set size set to 100 and 500, respectively; p-value threshold of .05; and Benjamini-Hochberg adjustment for multiple comparisons. Enriched terms were plotted using ggplot2 (version 4.0.0) (Wickham, 2016). Synaptic gene ontology (SynGO) analysis (release 1.2) (Koopmans et al., 2019) was used to identify and plot synaptic proteins and associated biological processes, specifically. All significantly up- and downregulated proteins were separately included in the analysis, with all annotations and requiring at least three protein counts to be included. Network maps of the top two enriched SynGO terms and daughter terms for up- and downregulated proteins were visualized using Microsoft PowerPoint 2024.

### Immunocytochemistry and imaging

To characterize cell identities and quantify localized proteins of interest, networks grown on 8-well chamber slides were fixed at 52 and 56 DIV. After washing the cells once using Dulbecco’s Phosphate Buffered Saline with calcium and magnesium (PBS; Sigma-Aldrich, D8662), networks were fixed for 15 min in 3% glyoxal solution (Richter et al., 2018) consisting of 71% distilled water, 20% ethanol absolute (Kemetyl, 2005786), 8% glyoxal (Sigma-Aldrich, 128465) and 1% acetic acid (Sigma-Aldrich, 1.00063). The glyoxal solution was removed before the cells were washed twice with PBS for 5 min and stored at 4°C until primary antibody staining. After the PBS was removed, the cells were permeabilized in 0.5% Triton-X (Sigma-Aldrich, 1.08643) for 5 min, followed by two 5 min washes using PBS. Blocking was subsequently performed by incubation in 5% goat serum (Sigma-Aldrich, G9023) for 30 min at RT on a rotating platform. The blocking solution was replaced by primary antibodies diluted in 5% goat serum in PBS. All primary and secondary antibodies that were used are listed in Table 2. Cells with the primary antibodies were incubated overnight on a tilting platform at 4°C. The next day, the primary antibodies were removed and the cells were washed three times with PBS for 5 min. The PBS was removed and replaced by secondary antibodies diluted to a 1:500 concentration in 5% goat serum in PBS. Cells were incubated with secondary antibodies for 3h at RT on a rotating platform. Secondary antibodies were removed and replaced with 8 uM Hoechst in PBS for 10 min. Hoechst was removed before the cells were washed twice using PBS, and one final time using distilled water. Cover glasses were mounted onto the cells before storage at 4°C until imaging using an EVOS M5000 microscope (Invitrogen) with a 20x/0,75 NA Olympus AMEP4734 objective, and LED light cubes DAPI (AMEP4650), GFP (AMEP4651), TX-Red (AMEP4655), and CY5 (AMEP4656). Image pro-cessing was performed in Fiji v1.54r.

**Table 2.**
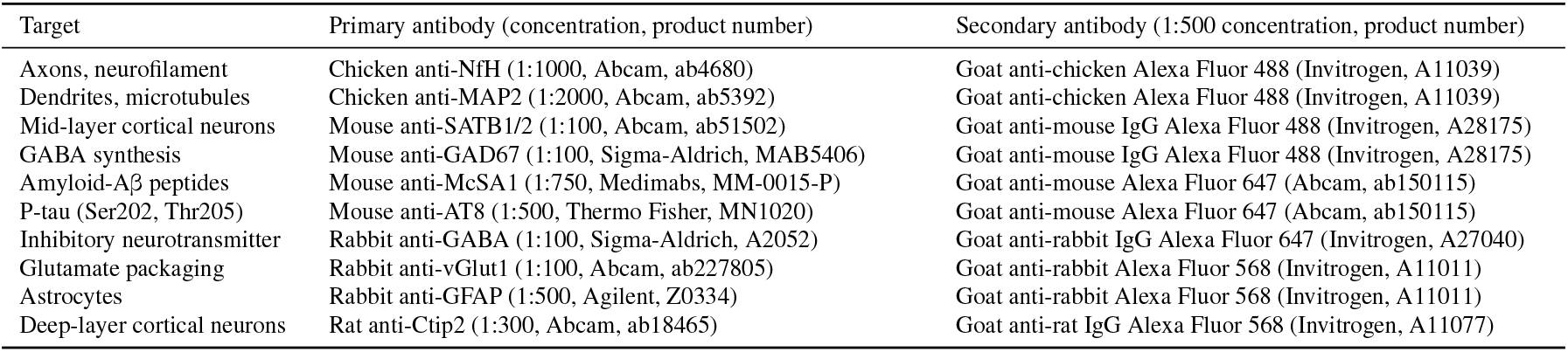
Primary and secondary antibodies used.

### Neurite quantification

To assess neurite growth at 52 and 56 DIV, images were taken from three different wells per condition (group* time point) using an automated scan protocol with an EVOS M7000 microscope (Invitrogen) with a 10x/0,3 NA EVOS AMEP4981 objective. 75% of each well were automatically imaged from the center using autofocus for each field. Images were visually assessed and excluded from the analysis if being out of focus, containing large debris, or displaying cells along the coverslip walls. This resulted in a total of 151 (APOE3 52 DIV), 95 (APOE3 56 DIV), 95 (APOE4, 52 DIV), and 142 (APOE4 56 DIV) images per group to be included in the analysis. Neurite quantification was performed in Fiji v1.54r using the NeuroConnectivity macro set (Verschuuren et al., 2019) (Verstraelen et al., 2020) (Verstraelen et al., 2024) with neurite detection based on Pani et al. (2014). Kruskal-Wallis test was used for statistical comparisons because Shapiro-Wilk test indicated significant deviances from the normal distribution. Conover-Iman test with Bonferroni correction was used for post hoc pairwise comparisons. Statistical analysis and plotting were performed in RStudio (RStudio IDE 2025.09.2+418) (R Core Team, 2021) using packages readxl (version 1.4.5) (Wickham and Bryan, 2025), dplyr (version 1.2.0) (Wickham et al., 2025a), ggpubr (version 0.6.3) (Kassambara, 2025), DescTools (version 0.99.60) (Signorell, 2025), and tidyr (version 1.3.2) (Wickham et al., 2025c).

### Quantification of excitatory and inhibitory markers

Glutamatergic and GABAergic activity was assessed by immunolabeling of VGLUT1 and GAD67 at 52 DIV, following a methodology adapted from Nadadhur et al. (2017) and Nieweg et al. (2015). VGLUT1 is a transporter protein responsible for packaging glutamate into synaptic vesicles. GAD67 is a cytoplasmic enzyme involved in GABA synthesis. Images were taken from three different wells per group using an automated scan protocol with an EVOS M7000 microscope (Invitrogen) with a 20x/0.45 NA EVOS LWD AMEP4982 objective. 40% of each well were automatically imaged from the center using autofocus for each field. Images were visually assessed and excluded from the analysis as described above. This resulted in a total of 338 (APOE3) and 109 (APOE4) images per group to be included in the analysis. The densities of VGLUT1 and GAD67 were quantified using the cell density counting workflow in Ilastik (version 1.4.0) (Berg et al., 2019) (Sommer et al., 2011) and normalized by MAP2 area. Because Levene’s test indicated that the assumption of equal variances was not met, groups were compared using the Welch t-test. Statistical comparisons and plotting were performed in RStuido (RStudio IDE 2025.09.2+418) (R Core Team, 2021) using packages dplyr (version 1.2.0) (Wickham et al., 2025a), ggpubr (version 0.6.3) (Kassambara, 2025), and haven (version 2.5.5) (Wickham et al., 2025b).

## Results

### Aberrant dendrite morphology and altered excitatory and inhibitory balance in APOE4 networks

Immunocytochemistry at 52 DIV revealed the successful differentiation of excitatory and inhibitory cortical neurons and astrocytes (Fig. 2A-H). Cells in both APOE3 and APOE4 networks expressed the cytoskeletal proteins microtubule-associated protein 2 (MAP2) and neurofilament heavy chain (NfH) (Fig. 2A-H), inhibitory neuron markers glutamate decarboxylase 1 (GAD67) and γ-aminobutyric acid (GABA) (Fig. 2A-D), excitatory neuron marker vesicular glutamate transporter 1 (VGLUT1) (Fig. 2A-B), cortical neuron markers COUP TF1-interacting protein 2 (CTIP2) and special AT-rich sequence binding protein 1 and 2 (SATB1+2) (Fig. 2E-F), and astrocyte marker glial fibrillary acidic protein (GFAP) (Fig. 2G-H).

**Fig. 2.**
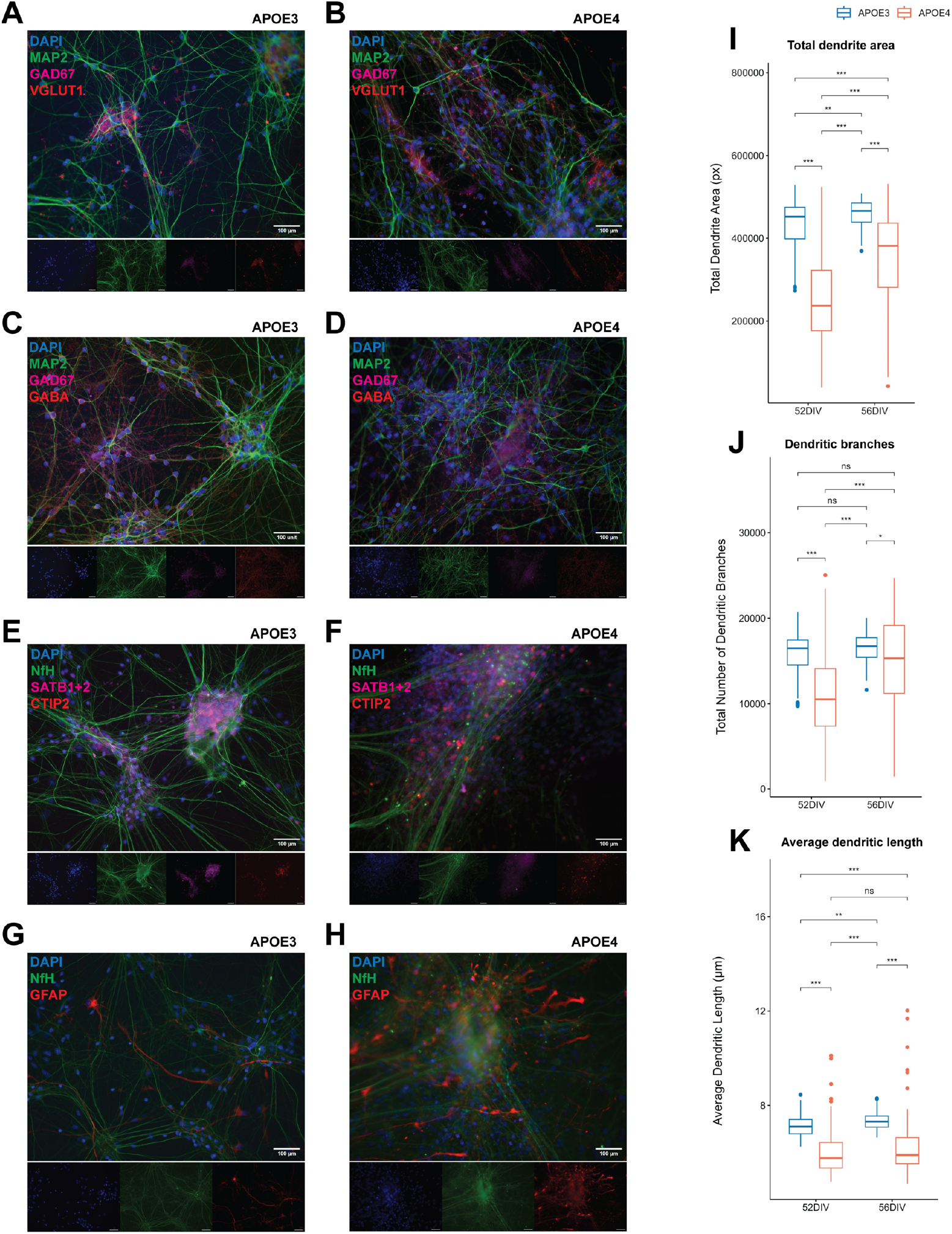
Left panel: Immunocytochemistry assays characterizing cell identity in APOE3 and APOE4 networks at 52 days in vitro (DIV). (**A-H**) The expression of MAP2 and NfH confirmed mature neuronal identity of cells in both groups. (**A-D**) Expression of excitatory and inhibitory neuron markers VGLUT1, GAD67, and GABA. (**E-F**) Expression of deep- and mid-layer cortical neuron markers CTIP2 and SATB1+2, respectively. (**G-H**) Expression of astrocyte marker GFAP. Scale bar: 100 um. Right panel: Quantification of dendrite morphology at 52 and 56 DIV. Box plots displaying differences between groups and time points in terms of (**I**) total dendrite area, (**J**) total number of dendritic branches, and (**K**) average dendritic length. Group legend is shared between figures. Kruskal–Wallis test followed by Conover-Iman test with Bonferroni correction for multiple comparisons. ***p<.001, **p<01, *p<.05, ns, not significant.

Quantification of dendrite morphology, indicated by fluorescent MAP2-labeling at 52 and 56 DIV, showed a significantly smaller total dendrite area in APOE4 neurons compared to APOE3 at both time points (Fig. 2I, p<.001). However, both groups showed an increase in total dendrite area over time, larger for APOE4 (p<.001) than for APOE3 (p<.01). In line with this, the total number of dendritic branches was smaller in APOE4 at both 52 DIV (p<.001) and 56 DIV (p<.05), with a significant time-dependent increase found only in APOE4 networks (Fig. 2J, p<.001). Lastly, the average dendritic length was shorter in APOE4 than APOE3 neurons at both time points of investigation (Fig. 2K, p<.001). Although a small, time-dependent increase in dendritic length was shown in APOE3 (p<.01), it remained stable in APOE4 networks.

Quantification of gluatamatergic and GABAergic neuronal activity by fluorescent VGLUT1 and GAD67 labeling at 52 DIV revealed an imbalance between excitatory and inhibitory synaptic transmission in APOE4 networks (Fig. 3A-B; Fig. 3G-H). Normalized to MAP2 area, APOE4 showed a slightly higher VGLUT1 density (Fig. 3G, p<.05) and a significantly lower GAD67 density than APOE3 neurons (Fig. 3H, p<.001).

**Fig. 3.**
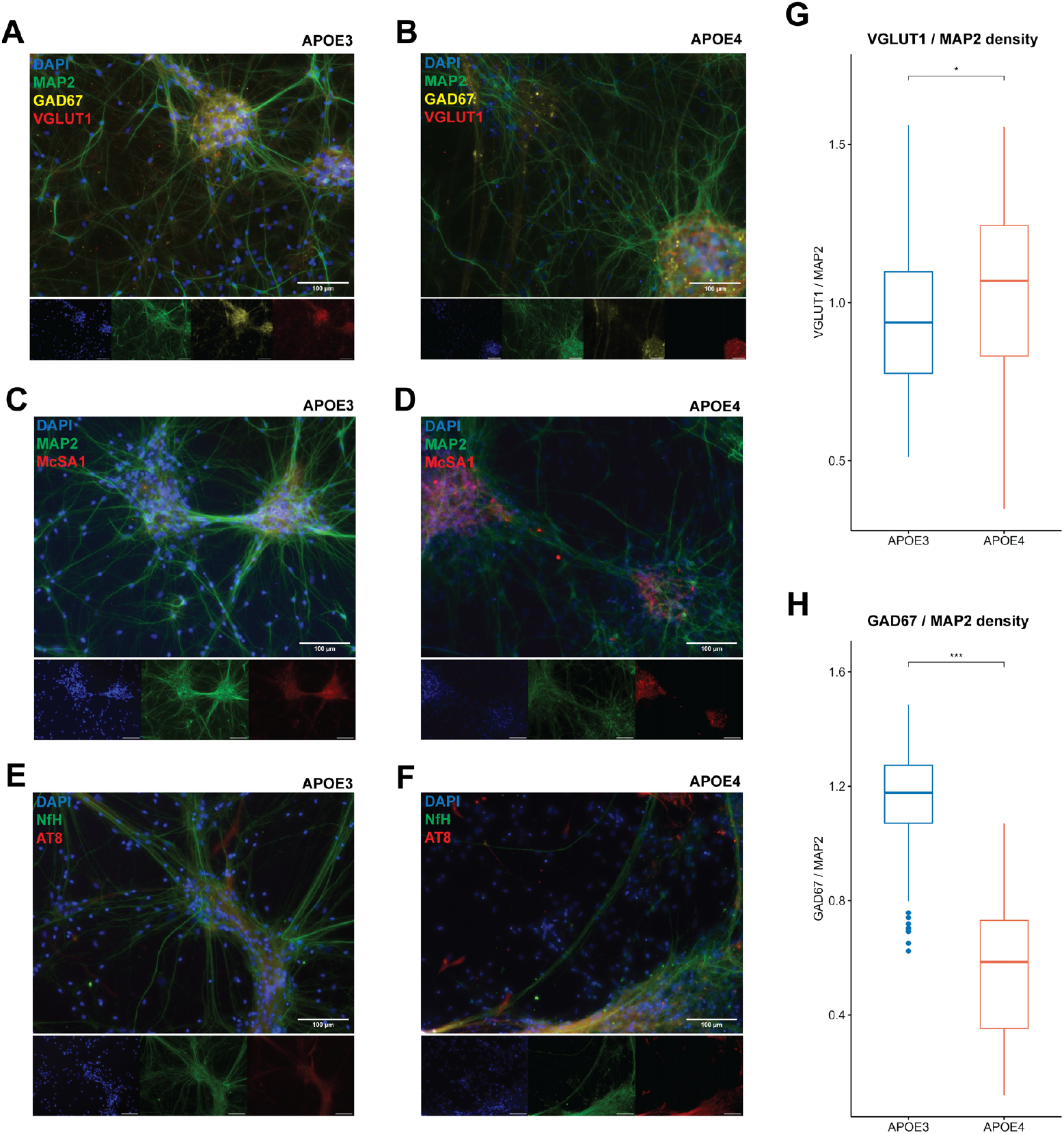
Left panel: Immunolabeling of APOE3 and APOE4 networks at 52 days in vitro (DIV). (**A-B**) Immunolabeling of excitatory and inhibitory neuron proteins VGLUT1 and GAD67, respectively, co-stained with neuronal marker MAP2. Scale bar: 100 um. Right panel: Quantification of excitatory and inhibitory immunomarkers at 52 DIV. (**G-H**) Quantification of VGLUT1 (top) and GAD67 (bottom), normalized by MAP2 density. Box plots displaying differences between groups. Welch t-test. ***p<.001, *p<.05.

In terms of Aβ and tau expression, McSA1- and AT8-positive cells were identified in both groups (Fig. 3C-F). McSA1, a marker for Aβ peptides, and AT8, labeling tau protein (Ser202/Thr205), are involved in both healthy and pathological functions, and are dynamically expressed during neurodevelopment (Bruno et al., 2025) (Weir et al., 2024) (Goedert et al., 1995). Aβ peptides are tightly regulated and serve healthy functions in neurons when expressed at normal levels (Zhang et al., 2024). Hyperphosphorylated tau (Ser202/Thr205) is typically considered pathological, but recent studies have also reported its abundance in young healthy neurons (Bruno et al., 2025) (Weir et al., 2024). Further complicating the analysis of the dynamic expression and function of these proteins, is that cultured neurons, even as part of the same network, may be in different pathological stages. Hence, to avoid introducing systematic statistical bias by regression towards the mean, the expressions of McSA1 and AT8 were not quantified.

### Reduced firing rate, synchrony, and transitivity in APOE4 networks, and increased inhibition and assortativity

Longitudinal microelectrode array recordings revealed a significantly reduced firing rate in APOE4 compared to APOE3 networks, with the strongest divergence apparent from 52 DIV and onward (Fig. 4A-B, p<.05). However, a larger fraction of the spikes were contained within electrode bursts in APOE4 networks (Fig. 4C-D, p<.05). Moreover, APOE4 networks displayed a larger fraction of inhibitory versus excitatory synaptic links at 70-82 DIV, with APOE4 showing approximately 40% compared to APOE3 networks with approximately 20% inhibitory connections (Fig. 4E-F, p<.001). The network synchrony, functional integration, and efficiency of information flow was hampered in APOE4 networks, indicated by no network bursting (Fig. 4G), and reduced global routing efficiency (Fig. 5A-B, p<.05). Network bursts were detected in APOE3 networks from 64 DIV (Fig. 4G). Both APOE4 and APOE3 networks possessed information processing capacity, shown by small world propensities (SWP) >.6 (Fig. 5C-D, p=.121) (Muldoon et al., 2016). While the SWP in APOE4 networks stabilized around 1.0 from 50 DIV, it decreased in APOE3 networks and stabilized around .6 from 64 DIV and throughout the experiment. APOE4 networks displayed a higher assortativity (Fig. 5E-F, p<.001), which measures the correlation between node degrees on two opposite ends of an edge. However, APOE4 networks showed lower transitivity than APOE3 networks (Fig. 5G-H, p<.01), which measures the ratio of triangles to triplets, i.e. parallel pathways to the same target nodes. Table 3 shows the fitted generalized linear mixed models for all electrophysiology metrics and both groups, with their estimated means, standard errors, confidence intervals, p-values, distributions, and link functions.

**Fig. 4.**
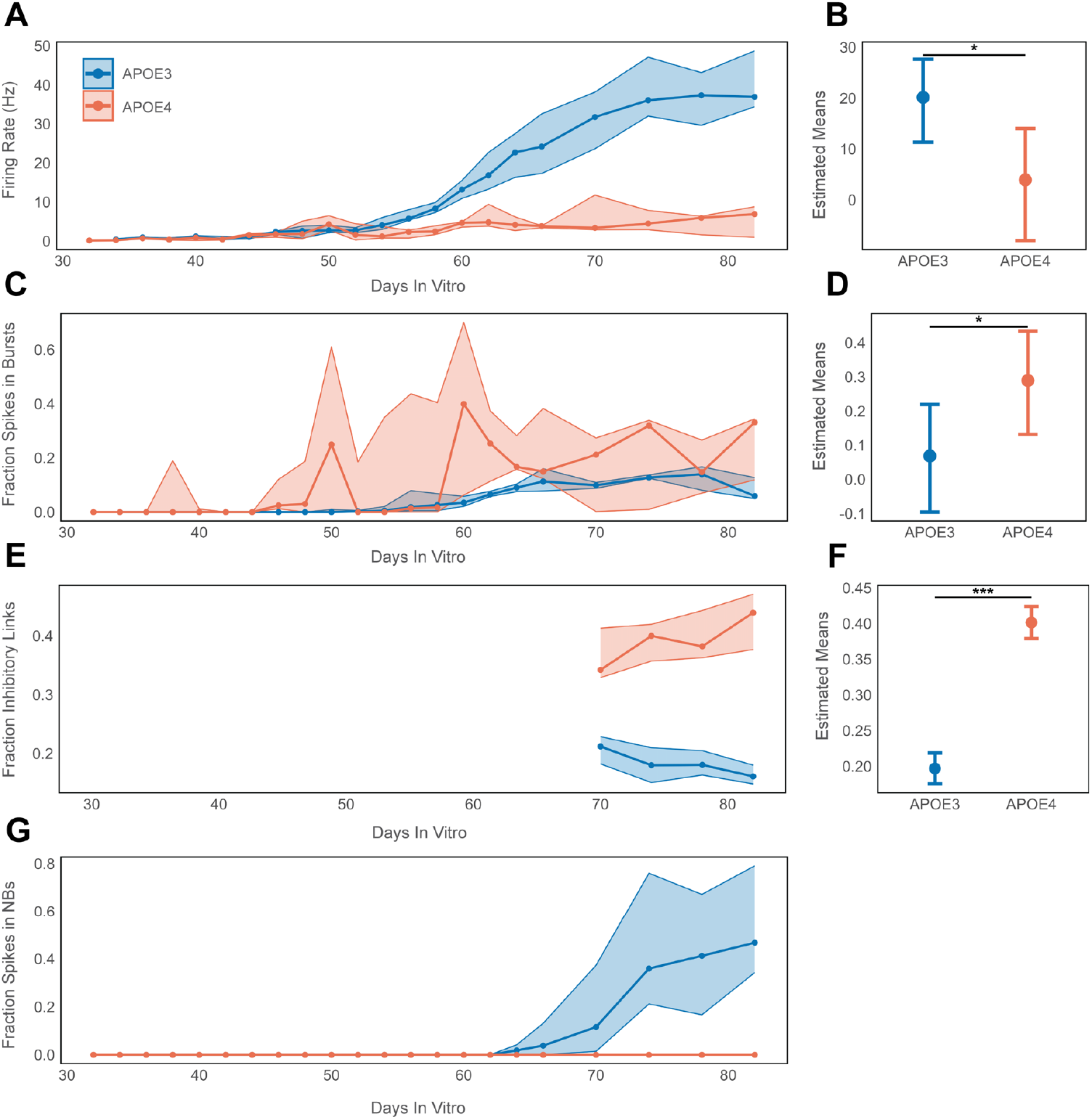
Network activity in APOE3 and APOE4 networks. (**A-B**) Firing rate in Hz, (**C-D**) fraction of spikes in bursts, (**E-F**) fraction of inhibitory synaptic links, and (**G**) fraction of spikes in network bursts (NBs). Left panel: Neuronal network activity over time. Solid lines and circles illustrate median value for all networks in each group. Shaded areas show upper and lower quartile. Group legend is shared between figures. Right panel: Generalized linear mixed models of estimated means with whiskers showing 95% confidence interval for each electrophysiology metrics. ***p<.001, *p<.05.

**Fig. 5.**
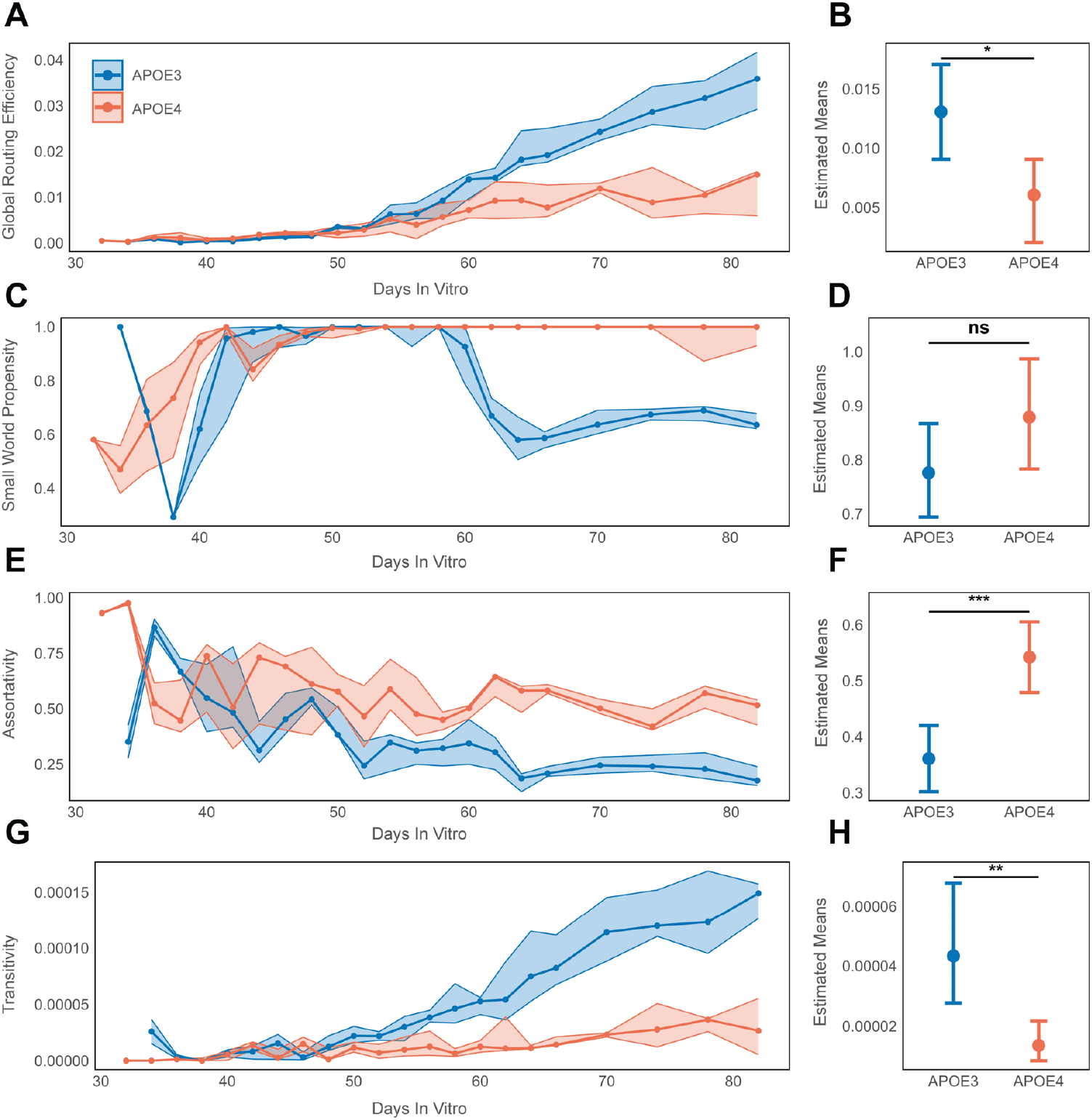
Network activity in APOE3 and APOE4 networks. (**A-B**) Global routing efficiency, (**C-D**) small world propensity, (**E-F**) assortativity, and (**G-H**) transitivity. Left panel: Neuronal network activity over time. Solid lines and circles illustrate median value for all networks in each group. Shaded areas show upper and lower quartile. Group legend is shared between figures. Right panel: Generalized linear mixed models of estimated means with whiskers showing 95% confidence interval for each electrophysiology metrics. ***p<.001,**p<01, *p<.05, ns, not significant.

**Table 3.** Estimated Means and Selected Target Distributions and Link Functions for the Electrophysiology Metrics.

| Group | Parameter | Estimated Means | SE | Lower 95% CI | Upper 95% CI | p-value<sup>a</sup> | Distribution (link function) |
| --- | --- | --- | --- | --- | --- | --- | --- |
| APOE3 | Firing Rate | 20,150 | 3,579 | 11,328 | 27,635 | <.05 | Gamma (log) |
| APOE4 | Firing Rate | 3,919 | 4,862 | -8,003 | 14,008 | <.05 | Gamma (log) |
| APOE3 | Fraction Spikes in Bursts | ,068 | ,071 | -,095 | ,219 | <.05 | Gamma (log) |
| APOE4 | Fraction Spikes in Bursts | ,289 | ,067 | ,131 | ,433 | <.05 | Gamma (log) |
| APOE3 | Fraction Inhibitory Links | ,197 | ,011 | ,175 | ,219 | <.001 | Normal (identity) |
| APOE4 | Fraction Inhibitory Links | ,401 | ,011 | ,379 | 424 | <.001 | Normal (identity) |
| APOE3 | Global Routing Efficiency | ,013 | ,002 | ,009 | ,017 | <.05 | Gamma (log) |
| APOE4 | Global Routing Efficiency | ,006 | ,002 | ,002 | ,009 | <.05 | Gamma (log) |
| APOE3 | Small World Propensity | ,776 | ,041 | ,694 | ,867 | ,121 | Gamma (log) |
| APOE4 | Small World Propensity | ,879 | ,048 | ,783 | ,987 | ,121 | Gamma (log) |
| APOE3 | Assortativity | ,361 | ,026 | ,302 | ,420 | <.001 | Normal (identity) |
| APOE4 | Assortativity | ,542 | ,028 | ,479 | ,605 | <.001 | Normal (identity) |
| APOE3 | Transitivity | 4,330 E-05 | 8,634 E-06 | 2,771 E-05 | 6,766 E-05 | <.01 | Gamma (log) |
| APOE4 | Transitivity | 1,363 E-05 | 2,872 E-06 | 8,546 E-06 | 2,173 E-05 | <.01 | Gamma (log) |

SE, standard error, CI, confidence interval.
<sup>a</sup> Bonferroni adjusted.

### Upregulation of synaptic and intracellular signaling, ion homeostasis, and dendritic processes in APOE4 neurons, and downregulation of cell adhesion, motility, and actin cytoskeleton

Enrichment analysis of differentially expressed synaptosome proteins (DEPs) showed dysregulation of synaptic signaling in APOE4 neurons, including upregulation of chemical synaptic transmission, anterograde trans-synaptic signaling, learning, glutamatergic synapse, neurotransmitter transport, synaptic and exocytic vesicle, receptor ligand activity, and signaling receptor activity (Fig. 6A-B). Proteins covered by the ontology terms dendrite and dendritic tree were also upregulated in APOE4 neurons (Fig. 6A), while actin-related terms were downregulated, including actin cytoskeleton, actin filament binding, and actin binding (Fig. 6C-D). Several terms involved in ion dynamics were upregulated in APOE4, including monoatomic and inorganic ion homeostasis, metal ion transmembrane transporter activity, and ion channel activity (Fig. 6B). APOE4 neurons also showed dysregulated intracellular signaling, in particular upregu-lation of G-protein coupled receptor signaling, and cyclic nucleotide phosphodiesterases (enzymes that break down the intracellular messengers cAMP and cGMP) (Fig. 6A). Several terms associated with cell adhesion and motility were downregulated in APOE4 neurons, including cell-cell adhesion, cell-substrate junction, integrin and cadherin binding, ruffle, and cell leading edge (Fig. 6C-D). Terms related to DNA transcription, vasculature, and transport were also dysregulated in APOE4 neurons (Fig. 6B-D). Lastly, several membrane-related terms were upregulated in APOE4, including postsynaptic membrane, synaptic vesicle membrane, and transport vesicle membrane (Fig. 6B)

**Fig. 6.**
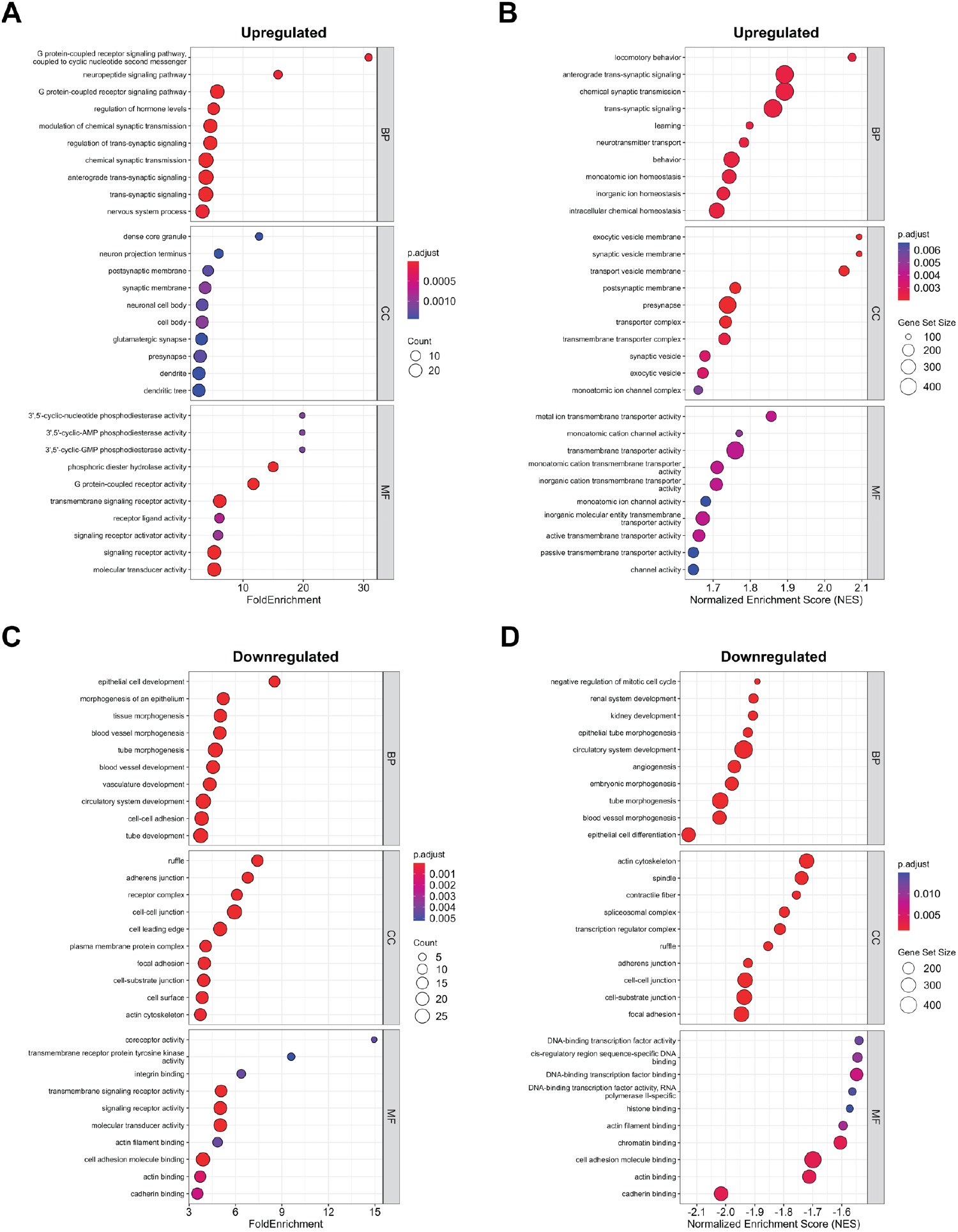
Enriched gene ontology (GO) terms for “Biological Process” (BP), “Cellular Component” (CC) and “Molecular Function” (MF) associated with differentially expressed proteins in APOE4 neurons. (**A**) Dot plot of the top ten upregulated terms with highest fold change in APOE4 neurons, identified by GO over-representation analysis (ORA). (**B**) Dot plot of the top ten upregulated terms with highest normalized enrichment scores (NES) in APOE4 neurons, identified by gene set enrichment analysis (GSEA). (**C**) Dot plot of the top ten downregulated terms with highest fold change in APOE4 neurons, identified by GO ORA. (**D**) Dot plot of the top ten downregulated terms with highest NES in APOE4 neurons, identified by GSEA. Adjusted p-values and protein counts are illustrated by dot color and dot size, respectively.

Synaptic gene ontology (SynGO) analysis of DEPs revealed upregulation of terms associated with synaptic signaling in APOE4 neurons, including chemical synaptic transmission, and retrograde trans-synaptic signaling by endocannabinoid (Fig. 7A-B). Synaptic organization and structure were also dysregulated in APOE4 neurons, with both up- and downregulated terms (Fig. 7A-D).

**Fig. 7.**
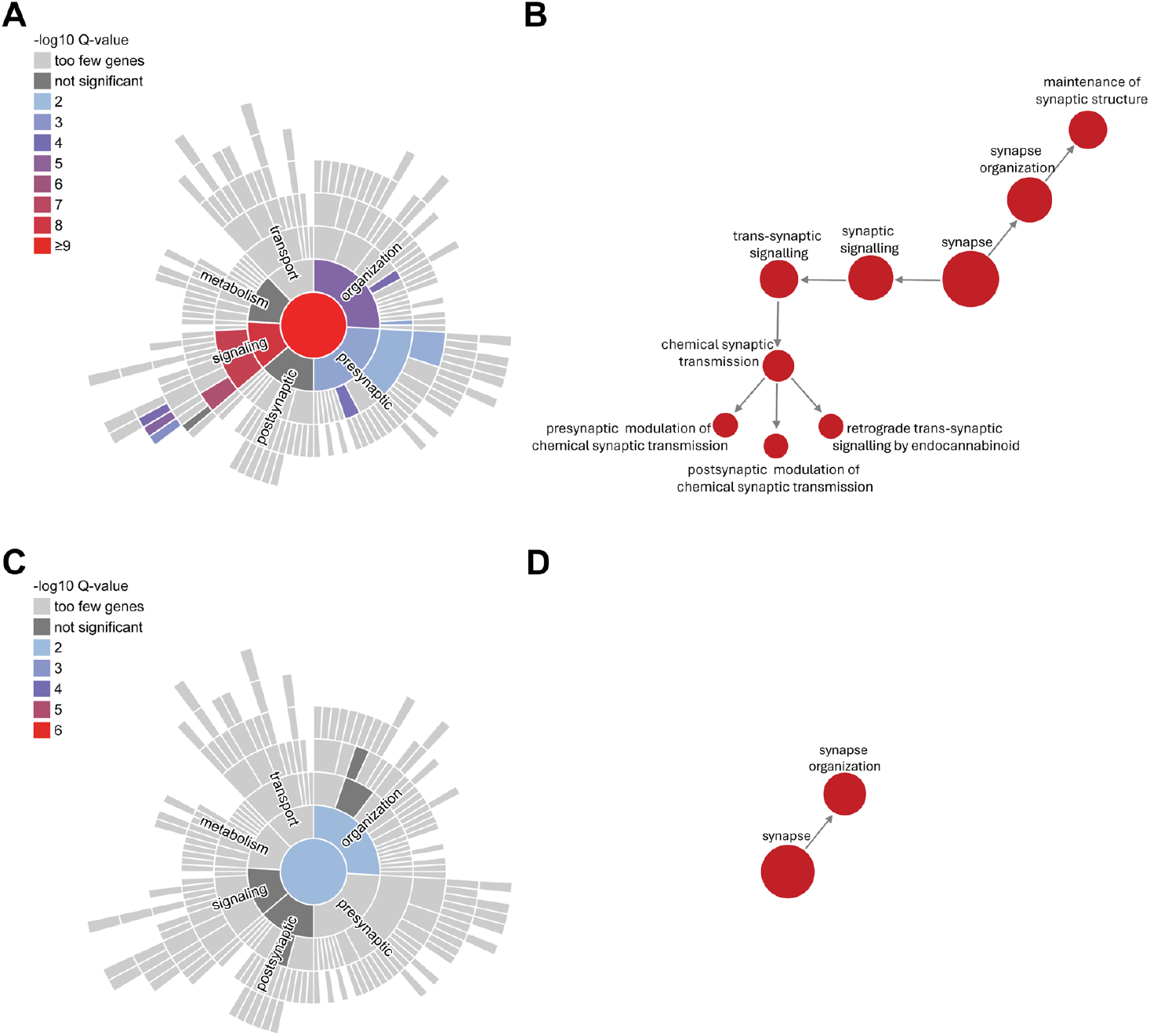
Enriched synaptic terms (left), and their associated enriched daughter terms (right) from the syntaptic gene ontology (SynGO) database. (**A-B**) Terms associated with upregulated differentially expressed proteins (DEPs) in APOE4. (**C-D**) Terms asociated with downregulated DEPs in APOE4.

238 unique DEPs were identified when comparing APOE4 and APOE3 neurons (Fig. 8A-B). DEPs with the highest log10 P-values included the Rho GTPase Activating Protein 36 (ARHGAP36), involved in neuronal development and signal transduction by inhibiting Protein Kinase A (Nam et al., 2019) (Eccles et al., 2016); Phosphodiesterase 10A (PDE10A), a dendritic spine-localized enzyme that regulates cyclic nucleotide signaling (Russwurm et al., 2015); Gonadotropin Releasing Hormone 1 (GNRH1), mostly known for its role in reproductive signaling, expressed in neurons which, in fish, are connected by gap junctions and facilitate synchronous firing of neuronal populations (Ma et al., 2015); Protein Phosphatase 1 Regulatory Inhibitor Subunit 1B (PPP1R1B), which encodes the DARPP-32 protein, involved in the integration of synaptic signals and regulation of the phosphorylation of neurotransmitter receptors and ion channels (Svenningsson et al., 2004); and Crystallin Mu (CRYM), which controls neurotransmitter release and is important for maintaining excitatory and inhibitory balance in neuronal networks (Ollivier et al., 2024).

**Fig. 8.**
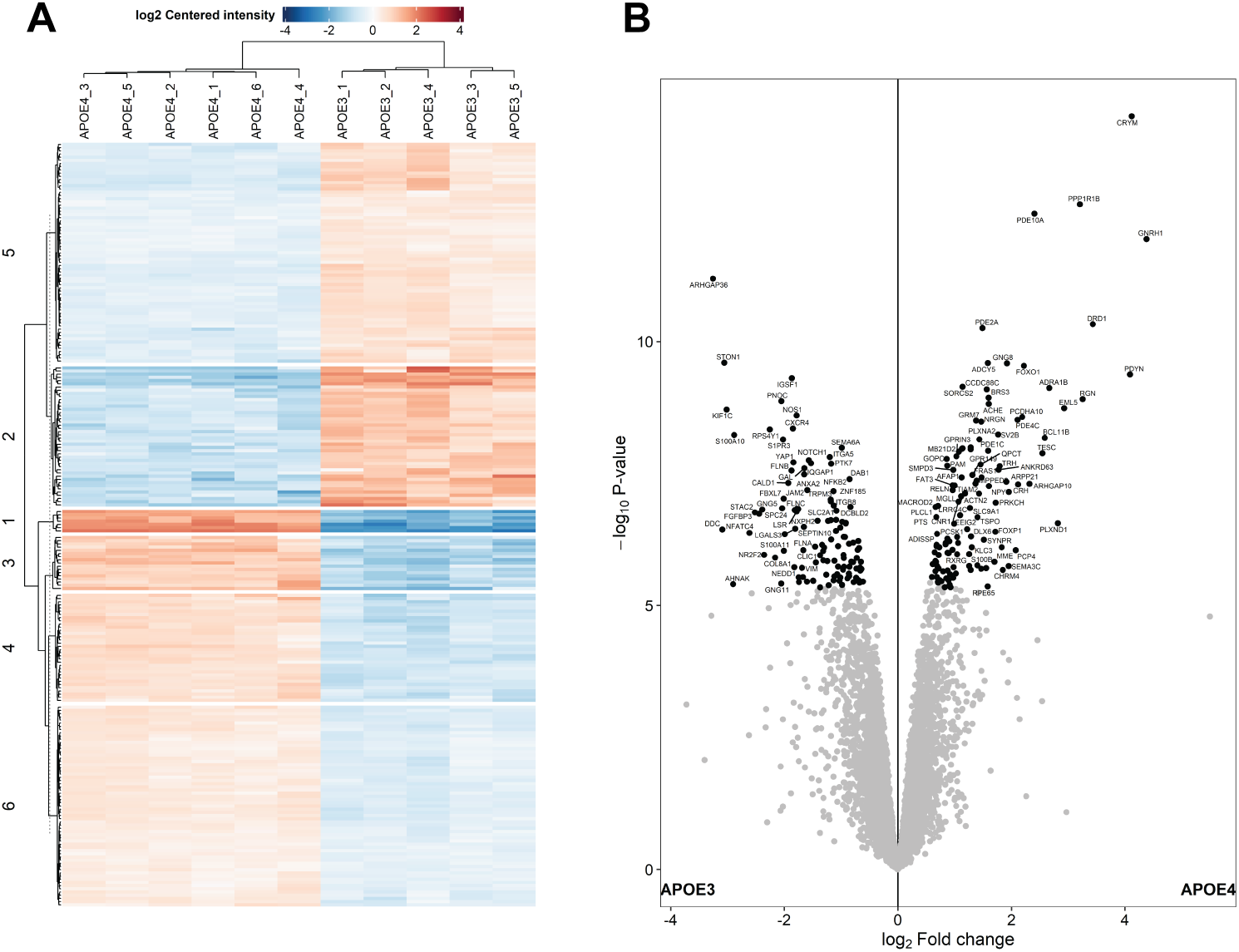
238 differentially expressed proteins (DEPs) were identified in isolated synaptosomes. (**A**) Heatmap and (**B**) volcano plot of the significant DEPs in APOE4 neurons compared to isogenic APOE3 control neurons.

## Discussion

In this study, we demonstrate that human neuronal networks with inherent vulnerability to Alzheimer’s disease (AD) by carrying homozygous APOE4, exhibit reduced firing rates, dynamic and time-dependent excitatory and inhibitory (E/I) imbalance, impaired synchronization, and reduced global information flow compared to APOE3 isogenic control networks. Topologically, APOE4 networks showed increased assortativity, which makes active pathways more resilient to random edge removal, albeit at the cost of rendering the networks susceptible to pathological spread. Furthermore, APOE4 displayed reduced transitivity, which impairs the functional flexibility of the networks. Underlying these network-level vulnerabilities, synaptosome proteomics revealed dysregulation of synaptic organization and signaling,ion dynamics, and intracellular signaling. Structurally, we observed reduced dendritic length, branching, and total dendrite area in APOE4 networks at 52 and 56 days in vitro (DIV), accompanied by dysregulation of actin, dendrite, and cell adhesion pathways. Together, these findings suggest that while APOE4 networks retain a baseline capacity for signal transmission and computation, the reliance on compensatory but fragile mechanisms renders them vulnerable to the additional pathological burden that arises with aging and disease progression.

Our proteomics results of dysregulated synaptic and intracellular signaling map closely onto pathways that are repeatedly reported as early and central substrates of synaptic impairment and dysfunctional neuronal network dynamics in AD. We found dysregulation of chemical synaptic transmission, glutamatergic synapse, neurotransmitter transport, synaptic vesicle, postsynaptic membrane, receptor ligand activity, ion homeostasis, G-protein coupled receptor signaling, and cyclic nucleotide phosphodiesterases in APOE4 neurons (Fig. 6A-D) (Fig. 7A-D). Previous proteomic and meta-analytic studies show that presynaptic vesicle machinery, neurotransmitter release, and postsynaptic receptor complexes are among the earliest and most consistently altered processes in AD patient brains (Griffiths et al., 2025) (Yarbro et al., 2025) (de Wilde et al., 2016). By analyzing the synaptic proteosome of postmortem patient tissue, Griffiths et al. (2025) identified a progressive trajectory of synaptic impairment, starting with early metabolic and presynaptic dysfunction, followed by inhibitory synapse alterations in the middle stages of disease, before a late-stage selective vulnerability of glutamatergic synapses (Griffiths et al., 2025). The synaptic proteomic signature we identify in APOE4 neurons, spanning presynaptic vesicle machinery, postsynaptic receptor signaling, and ion homeostasis, thus recapitulates hallmarks of early-stage synaptic AD pathology.

Structurally, we observed reduced dendritic length and branching in APOE4 neurons (Fig. 2I-K). Cytoskeletal remodeling and neurite degeneration have been consistently reported in AD (Zhang et al., 2024) (Zhang et al., 2025) (Blanquie and Bradke, 2018) (Kao et al., 2010) (Salluzzo et al., 2023) (Yarbro et al., 2025) (de Wilde et al., 2016). The hyperphosphorylation and aggregation of tau into neurofibrillary tangles disrupt microtubule stability, leading to neurite dystrophy, spine loss, and impaired axonal transport (Zhang et al., 2025). Altered expression of cell adhesion molecules has also been strongly linked to disrupted neurite extension, abnormal axon bundling, and progressive degeneration in neurodegenerative diseases (Salluzzo et al., 2023) (Andersen et al., 2024). This corresponds well to our findings of dysregulated cell adhesion and motility in APOE4 networks, including cell-cell adhesion, cell-substrate junction, integrin and cadherin binding, ruffle, and cell leading edge (Fig. 6A-D). In AD and related tauopathies, shifts in tau isoform ratios and cell adhesion molecules signaling are proposed to shift the balance toward either excessive cytoskeletal stabilization or fragmentation (Salluzzo et al., 2023) (Andersen et al., 2024). In fact, during the progression of AD, degenerative (tangle formation and neurite beading) and regenerative (sprouting neurites with growth cones) processes have been observed simultaneously, suggesting that aberrant or mis-targeted activation of cytoskeletal growth programs in a pathological cellular environment leads to abnormal neurite growth rather than effective circuit repair (Kao et al., 2010) (Blanquie and Bradke, 2018). In light of this, our findings of dysregulated cell adhesion, dendritic trees, and actin binding, impaired dendritic growth and branching, dysregulated synaptic organization and signaling, and vulnerable and hypoactive electrophysiological dynamics in APOE4 networks, constitute a cohesive, multifactorial perspective on early AD pathology. The self-organization and growth of APOE4 networks are hampered, which could drive neurons to establish synaptic connections at suboptimal sites (Nwabuisi-Heath et al., 2014) (Jain et al., 2013), thus reducing electrophysiological efficiency. The latter is manifested in our data by aberrant synaptic signaling, shown in the electrophysiological results as reduced firing rate and global routing efficiency, and in the proteomics results as dysregulated neurotransmitter transport, synaptic and exocytic vesicle, receptor ligand activity, and signaling receptor activity. This is also in line with previous findings linking APOE4 to dysregulated formation (Mauch et al., 2001), function (Chen et al., 2025), and plasticity of synapses (Chen et al., 2010). Importantly, however, the temporal and causal relationships between these mechanisms remain unresolved. It is unclear whether structural defects precipitate synaptic instability or whether synaptic dysfunction triggers cytoskeletal remodeling.

The time-dependent reduced firing rates of APOE4 compared to APOE3 networks (Fig. 4A-B) were accompanied by dysregulated monoatomic inorganic cation dynamics in the proteomics data (Fig. 6A-D). These ions include sodium (Na^+^), potassium (K^+^), and calcium (Ca^2+^), which are involved in membrane potential maintenance, action potential generation, neurotransmitter release, and synaptic plasticity (Bhoi et al., 2025) (Lisman, 1997) (Csicsvari et al., 1998) (Kobayashi et al., 2024). Interestingly, APOE4 and APOE3 showed comparable firing rates up until 52 DIV, after which the spiking activity of APOE3 networks strongly increased and diverged from that of APOE4. At 52 DIV, APOE4 neurons showed increased VGLUT1/GAD67 ratios (Fig. 3G-H). As the electrophysiological trajectory of APOE3 involved increased activity from this time point, the upregulation of excitatory synaptic transmission in APOE4 networks may represent a means to preserve signal transmission when synaptic efficiency begins to fail, as indicated happening by the proteomics data (Fig. 7A-D). Previous studies have demonstrated that elevated VGLUT1 levels increases the probability of neurotransmission (Herman et al., 2014) (Taipala et al., 2022). However, the longitudinal electrophysiological data shows that the increased glutamate packaging in APOE4 neurons at 52 DIV is not sufficient to compensate for their impaired synaptic function, i.e. their firing rates remain lower than those of APOE3 networks after 52 DIV. Furthermore, increased glutamate packaging and release, without functional pathways for utilizing and handling this glutamate efficiently, can lead to excitotoxicity (Targa Dias Anastacio et al., 2022). Previous studies have reported impaired glutamate uptake and conversion by astrocytes (Targa Dias Anastacio et al., 2022). Increased levels of glutamate available in the synaptic cleft increases the activation of N-metyl-D-aspartat (NMDA) receptors, which can lead to excessive Ca^2+^ influx and thereby neurotoxicity (Targa Dias Anastacio et al., 2022). Aβ has been reported to accelerate these processes by reducing glutamate uptake, increasing NMDA activation of glutamate, and forming ion-permeable pores in the plasma membrane (Targa Dias Anastacio et al., 2022). Hence, the failed compensatory attempt of APOE4 to increase spiking by elevating glutamate levels at 52 DIV, could explain the dysregulated ion homeostasis and intracellular signaling identified at 64 DIV (Fig. 6A-D). However, between 70-82 DIV, analysis of E/I links indicates that the early elevated excitatory drive of APOE4 networks had diminished and shifted to increased inhibition. Based on previous reports of glutamate excitotoxicity and selective vulnerability of glutamatergic neurons, our findings of increased fraction of inhibitory links at late time points could therefore either suggests damage and dysfunction of excitatory neurons (Leng et al., 2021), or adaptive homeostatic plasticity where the networks upregulate inhibition to counteract glutamate excitotoxicity (Muir et al., 1996) (Bayón-Cordero et al., 2022). Nevertheless, APOE4 networks display a dynamic temporal shift in the E/I balance, closely resembling electrophysiological signatures reported during Alzheimer’s disease progression in patients and animals. Hyperexcitability is often observed at the early stages of the disease, before transitioning to hypoexcitability as the disease progresses (Burns et al., 2026) (Li et al., 2025) (Palop and Mucke, 2010) (Castanho et al., 2025) (Zerbi et al., 2014) (Fadel et al., 2025) (Targa Dias Anastacio et al., 2022). Very few in vitro have previously monitored network activity as extensively and over long enough time courses to identify similar transitions (Lin et al., 2018) (Ghatak et al., 2024) (Targa Dias Anastacio et al., 2022). Furthermore, our findings of dynamic changes in synaptic function and resulting network activity, where both adaptive and maladaptive plasticity mechanisms are at play, mimics clinical reports of AD progression as a non-linear process (Popescu et al., 2020) (Jamalian et al., 2023) (Lim et al., 2022).

Importantly, both groups possessed information processing capacity, as suggested by small world propensities (SWP) >.6 (Fig. 5C-D) (Muldoon et al., 2016). The increased fraction of inhibitory links in APOE4 networks (Fig. 4E-F) likely contributes to sustain this computational capacity, as increased inhibition, when properly balanced with excitatory drive, stabilizes network dynamics (Bhoi et al., 2025), sharpens tuning (Willmore et al., 2011), and reduces noise (Wehr and Zador, 2003). This leads to more precise spikes and bursts that carry more information despite an overall lower firing rate (Sengupta et al., 2013). However, the absence of network bursting (Fig. 4G) and the hampered global routing efficiency in APOE4 networks from 54 DIV (Fig. 5A-B) indicate that APOE4 networks have not reached a coherent functional organization capable of fully efficient integration and distributed processing (Bauer et al., 2022) (Cabrera-García et al., 2021). The SWP of APOE3 networks stabilized around .6 from 64 DIV and through all later time points. For APOE4 networks, however, the SWP remained stable at 1 from 50 DIV and throughout the experiment. The repeated observations of SWP = 1 in APOE4 are due to the effects of bounding between 0 and 1. In the calculation of the SWP, if network clustering is greater than that observed in a comparable lattice network, or the characteristic path length is shorter than that observed in a comparable random network, the deviations from the null models are set to zero (Muldoon et al., 2016). When both conditions are present in the same network, this results in SWP = 1, and indicates that the network has strong small world characteristics. Although the APOE4 networks showed a higher SWP than APOE3 networks, this does not necessarily mean that these networks exhibit a healthier and more functional network architecture. Instead, the organization of APOE3 networks likely represents a more sustainable state, where homeostatic plasticity balances computational efficiency and wiring cost. After demonstrating excessive SWP associated with a metabolically active core zone in Parkinson’s disease networks, Ko et al. (2018) concluded that under pathological conditions, elevated SWP increases the metabolic cost of computation at critical hub nodes. In other words, maintenance of important pathways for information transfer increases the wiring cost. This is in line with our own previous findings of increased metabolic demands in Parkinson’s and ALS networks (Valderhaug et al., 2024) (Fiskum et al., 2026), and previous reports from human APOE4 networks (Budny et al., 2024). The increased assortativity of APOE4 networks in the present study (Fig. 5E-F) indicates that highly connected nodes and pathways are indeed prioritized and protected from perturbations, because nodes tend to connect to other nodes of similar connectivity strength. Previous studies have demonstrated that assortative networks are more resilient to random damage or edge removal, however, at the cost of rendering networks more susceptible to pathological spread (Newman, 2003) (Vazquez and Moreno, 2003) (Zhou et al., 2012) (Chen et al., 2023). Despite this risk, the higher assortativity of APOE4 networks can be understood as a compensatory mechanism to counteract the lower ratio of triangles to triples, transitivity, in the same networks (Fig. 5G-H), which entails an overall vulnerability to network failure, with few alternative routes and hence less flexibility when comparing all pairs of nodes. The metabolic vulnerability implied by elevated wiring costs and assortative topology is further reinforced by our proteomics findings of dysregulated ion dynamics in APOE4 neurons.

Maintaining ion gradients across neuronal membranes is among the most energy-demanding processes in the brain, with neurotransmission-related processes powered by Na^+^ pumping estimated to consume at least 55% of the total brain energy demand (Engl and Attwell, 2015). If AD pathophysiology is in part driven by a progressive asymmetry between energy demand and metabolic supply, as it has been proposed (Budny et al., 2024) (Ardanaz et al., 2022), it is plausible that one of the most energy-intensive processes in neurons would be among the first to exhibit proteomic dysregulation. Impaired active transport of Na^+^ and K^+^ would not only compromise the restoration of the resting membrane potential following action potential firing, but would also alter the intra- and extracellular ion concentrations that underlie single-neuron excitability and synaptic integration, ultimately feeding back onto the network-level instabilities in E/I balance and synchronization observed here. In light of this, ionic dysregulation may not merely represent a molecular correlate of network dysfunction, but an active and early contributor to the progressive functional deterioration that APOE4 networks appear predisposed to undergo.

At the same time point as the SWP in APOE3 declined,i.e. at 64 DIV, synchronized bursting was detected in these networks (Fig. 4G). Although network bursts contribute to distributed and synchronized processing, this can come at the cost of limiting fine-tuned information processing (Kobayashi et al., 2024) (Houben et al., 2025) (Gast et al., 2020). Houben et al. (2025) suggested that this happens due to synaptic resource depletion following network bursting, which constrains fine-grained temporal coding and thus limits information transmission. Kobayashi et al. (2024) showed that when stimuli trigger network bursts, the responses of individual neurons converge to stereotyped patterns dominated by the network state, thus providing a mechanism for network-burst dependent information loss. By maintaining network dynamics that keeps the SWP at the minimum required for computation, and engaging in network bursting, APOE3 networks save metabolic resources and mobilizes larger regions of the circuit for information processing, which in the long run is a more sustainable way of functioning while maintaining homeostasis.

The absence of network bursting in APOE4 networks at 64 DIV likely reflects a convergence of disruptions across multiple levels of neural circuit organization. Oscillating activity in neuronal networks depends on single-neuron excitability and E/I balance within recurrent circuits (Aradi and Maccaferri, 2004) (Fröhlich et al., 2006) (Blankenship and Feller, 2010) (Takahashi et al., 2010) (Turrigiano, 2011) (Penn et al., 2016), and both of these prerequisites were compromised in APOE4. Single-neuron excitability is itself shaped by intrinsic membrane properties, ion concentrations, and synaptic connectivity, as subthreshold dendritic signals are integrated to generate action potentials (Penn et al., 2016). Hence, both the dysregulated ion homeostasis and the reduced dendritic length, branching, and total dendrite area observed in APOE4 networks can be expected to disrupt the networks’ integration capacity. The dysregulation of synaptic signaling and organization pathways further erodes the recurrent connectivity on which fast network oscillations depend, while disruption of intracellular signaling cascades may impair homeostatic compensatory mechanisms to restore excitability (Rátkai et al., 2021). Finally, given that slow rhythmic activity depends on astrocyte-mediated neurotransmitter recycling (Huang et al., 2017), any APOE4-related astrocyte dysfunction in our co-culture networks could further suppress the emergence of even low-frequency coordinated activity. Together, this suggest that APOE4 disrupts network bursting not through a single failure point, but through a structural and molecular re-configuration that renders the networks unable to meet the interdependent cellular and synaptic conditions required synchronized network bursting.

In summary, our findings provide a multiscale characterization of how APOE4 reorganizes the structural, molecular, and functional architecture of human neuronal networks, establishing a mechanistic framework for early AD vulnerability. The combined effects of reduced dendritic complexity, dysregulated synaptic signaling, altered excitatory and inhibitory balance, and reconfigured network topology indicate that APOE4 networks operate in a constrained and metabolically demanding regime. Although baseline information processing capacity is preserved, the reduced redundancy and impaired integration observed here suggest limited flexibility in the face of additional stressors. Rather than simply diminishing activity, APOE4 appears to reorganize neuronal networks into a state that is compensatory in the short term but progressively susceptible to destabilization. These results underscore the importance of early interventions targeting synaptic organization, ion homeostasis, and circuit-level balance before network fragility becomes entrenched. More broadly, this study demonstrates the value of integrating structural, proteomic, and network-level analyses to elucidate how genetic risk factors translate into system-level dysfunction.

## AUTHOR CONTRIBUTIONS

**MBG:** Conceptualization, Methodology, Software, Formal Analysis, Investigation, Data Curation, Writing - Original Draft, Writing - Review & Editing, Visualization. **VF:** Methodology, Software, Formal Analysis, Investigation, Writing - Original Draft, Writing - Review & Editing. **AS:** Conceptualization, Resources, Writing - Review & Editing, Funding acquisition. **IS:** Conceptualization, Resources, Writing - Review & Editing, Supervision, Funding acquisition.

## ACKNOWLEDGEMENTS

Mass spectrometry-based analyses were performed by the Proteomics and Modomics Experimental Core (PROMEC), Norwegian University of Science and Technology (NTNU) and The Central Norway Regional Health Authority. This facility is a member of the National Network of Advanced Proteomics Infrastructure (NAPI), which is funded by the Research Council of Norway INFRASTRUKTUR-program (project number: 295910). Data storage and handling is supported under the NIRD/Notur project NN9036K.

